# Cardiomyocyte mechanical activity counteracts intraluminal calcium depletion in the transverse-axial tubular system during fast electrical stimulation

**DOI:** 10.64898/2026.01.09.698373

**Authors:** S Orós-Rodrigo, J Fu, J Greiner, J Madl, M Linder, CM Zgierski-Johnston, A Loewe, P Kohl, EA Rog-Zielinska

**Affiliations:** Institute for Experimental Cardiovascular Medicine, University Heart Center and Faculty of Medicine, University of Freiburg, Freiburg, Germany; Centre for Integrative Signalling Studies (CIBSS), University of Freiburg, Freiburg, Germany; Institute of Biomedical Engineering, Karlsruhe Institute of Technology (KIT), Karlsruhe, Germany

## Abstract

The transverse-axial tubular system (TATS) enables close structural and functional coupling between plasma membrane and sarcoplasmic reticulum of cardiomyocytes. It supports fast and efficient Ca^2+^-induced Ca^2+^ release upon cell depolarisation, crucial for excitation–contraction coupling in the heart. Due to the small diameter and tortuosity of individual tubules, the TATS forms a domain of restricted diffusive transport. It has previously been suggested that, as a consequence of an uneven distribution of Ca^2+^ influx and efflux pathways in TATS compared to outer surface plasma membrane domains of cardiomyocytes, cyclic electrical activity may lead to a gradual depletion of Ca^2+^ in the TATS. Here, we show experimentally that in mechanically uncoupled rabbit ventricular cardiomyocytes, electrical stimulation does indeed lead to an L-type Ca^2+^ channel-dependent gradual depletion of Ca^2+^ inside TATS, an effect that scales with pacing frequency. Ca^2+^ depletion was absent in freely contracting cardiomyocytes, presumably as a result of cyclic TATS deformation during cell shortening. This squeezes transverse TATS tubules and adds an advective contribution to, and thereby accelerates the, intra-TATS content exchange with bulk extracellular fluid. Our results reveal a novel mechanism of cardiac mechano-dependent auto-regulation, where the increased propensity for development of intra-TATS Ca^2+^ gradients at high electrical stimulation rates is mitigated by the coinciding mechanically induced TATS deformation, twice on each cycle in the heart (during diastolic stretch and systolic shortening), which accelerates luminal content exchange. Our study provides first insight into a novel facet of cardiac mechano-biology, whose auto-regulatory benefit may be reduced by TATS remodelling in disease.

## Introduction

Efficient contraction of the heart relies on the synchronous shortening of individual cardiomyocytes, which in turn is dependent on the electrical signal reaching all sites of Ca^2+^–induced Ca^2+^ release of the cell near-instantaneously. This is enabled by the presence of the transverse–axial tubular system (TATS). TATS forms an elaborate network of interconnected surface membrane invaginations with diameters between 50–600 nm, and it is composed of transverse and axial tubules (TT and AT, chiefly oriented either perpendicular or parallel to the long axis of the cardiomyocyte, respectively). A TATS is found both in ventricular and in atrial cardiomyocytes, although with subtle differences in structure and density. Pathological remodelling is associated with profound changes in TATS structure and function, including decreased density and regularity of TT and reduced coupling to intracellular Ca^2+^ stores.^1–5^

While TATS membranes are physically and electrically continuous with the ‘outer’ surface membrane, jointly forming the sarcolemma, its luminal content is continuous with bulk extracellular fluid. As a result of the extensive presence of TATS, no part of a healthy cardiomyocyte’s cytosol is further than ∼1 μm away from the sarcolemma, separating cell in-and outside. In the process of excitation–contraction coupling, depolarisation triggers Ca^2+^ influx from the extracellular space, both at the cell’s outer surface and inside TATS, through voltage-gated L-type Ca^2+^ channels (LTCC); this in turn triggers a much larger Ca^2+^ release from the sarcoplasmic reticulum, giving rise to the cytosolic Ca^2+^ transient that enables contraction.^6^

In steady-state, an amount of Ca^2+^ equal to what entered the cell from outside must be extruded back to the extracellular space, chiefly via the Na^+^/Ca^2+^ exchanger (NCX). The distribution of trans-sarcolemmal Ca^2+^ influx (LTCC) and Ca^2+^ efflux (NCX) pathways appears to be inhomogeneous between TATS and outer surface membranes. Previous reports suggest that 60–80% of LTCC current flows across TATS membranes, whereas NCX shows a less pronounced heterogeneity with only 40–70% located in TATS.^7–10^ As a result, Ca^2+^ influx from TATS into the cardiomyocyte during excitation may exceed Ca^2+^ extrusion from the cytosol back into the TATS lumen upon relaxation. This, together with restricted diffusion, estimated to be up to an order of magnitude slower in TATS than in bulk extracellular space (due to tortuosity and small diameter of tubules), could give rise to gradients in intraluminal TATS Ca^2+^ concentrations ([Ca^2+^]_TATS_) that differ from Ca^2+^ concentrations in the bulk (outer) extracellular space ([Ca^2+^]_o_).^4,7,11–15^ This could manifest itself in gradual intraluminal TATS Ca^2+^ depletion at high pacing rates, in particular in TATS elements that are more distant from the cell surface.

While the development of such ion gradients within TATS has not been directly demonstrated experimentally, previous studies, conducted using double-barrelled Ca^2+^-selective microelectrodes, have shown that extracellular Ca^2+^ concentration in the proximity of rabbit papillary muscles decreases by ∼2–3% during each beat, presumably as a consequence of Ca^2+^ uptake by the cell.^16–21^ Computational modelling studies support the quantitative plausibility of TATS Ca^2+^ depletion.^22,23^ While existence, and possible functional consequences, of TATS Ca^2+^ depletion remain unexplored, it is likely that – if present – they would have detrimental effects on excitation–contraction coupling.

Previously, we demonstrated cyclic beat-by-beat TT and AT deformations, giving rise to an advective contribution to TATS content exchange in cyclically contracting or stretched cardiomyocytes, with different dynamics in AT and TT (accelerated content exchange in both during stretch, and in TT, but not AT, during contractions). We furthermore found that the contraction-induced acceleration of TT luminal content exchange was related to contraction frequency and amplitude, suggesting that TT pumping is a ‘use-dependent’ process that scales with changes in heart rate and/or stroke volume.^24–26^ We therefore postulate that TT pumping could stabilise [Ca^2+^]_TATS_, representing a novel mechanism of cardiac auto-regulation, whereby increased propensity for TATS luminal concentration imbalances at high stimulation rates would be countered by a matching elevation in the number both of stretch- and of contraction-driven advection-cycles of TATS content (two pumping cycles per each electrical activation). Here, we performed measurements of [Ca^2+^]_TATS_ dynamics in mechanically uncoupled and in contracting rabbit cardiomyocytes, using a non-membrane-permeable fluorescent Ca^2+^ indicator and confocal imaging. We demonstrate that in mechanically uncoupled cells, electrical activity leads to an LTCC-dependent Ca^2+^ depletion in TT, but not AT. This effect scales with stimulation frequency, and – based on computational modelling – is expected to be reduce intracellular Ca^2+^ transients and tension development. TATS Ca^2+^ depletion can be prevented by allowing cells to contract. We propose that an uneven distribution of Ca^2+^ influx and efflux pathways across TATS and outer surface membrane leads to progressive Ca^2+^ depletion in TATS during electrical activity, an effect that is countered in mechanical active cardiomyocyte by TT pumping, driving advection-assisted TATS luminal content exchange.

## METHODS

### Animals

All investigations reported in this manuscript conformed to German animal welfare laws (TierSchG and TierSchVersV), were compatible with the guidelines stated in Directive 2010 / 63 of the European Parliament on the protection of animals used for scientific purposes, and were approved by the local authorities (Regierungspräsidium Freiburg, X-21 / 01R). New Zealand white rabbits (age ∼2 months, male and female, weight ∼2–2.5 g) were anesthetised using esketamine hydrochloride (Ketanest 25 mg/mL, 0.5 mL/kg body weight; Pfizer, New York, NY, USA) and xylazine hydrochloride (Rompun 2%, 0.2 mL/kg body weight; Bayer Vital, Leverkusen, Germany) injected intramuscularly. Euthanasia was induced by intravenous injection of sodium thiopental (25 mg/mL; Inresa Arzneimittel, Freiburg, Germany). Hearts were Langendorff-perfused with solution ‘A’, containing (in mM): 137 NaCl, 4 KCl, 1.8 CaCl_2_; 1 MgCl_2_, 10 HEPES, 10 glucose, 20 taurine, 10 creatine, 5 adenosine, 2 L-carnitine, pH 7.3 when oxygenated at 37°C, ∼330 mOsm; followed by solution ‘B’ containing (in mM): 138 NaCl, 0.33 NaH_2_PO_4_, 5.4 KCl, 0.5 CaCl_2_, 2 MgCl_2_, 10 HEPES, 10 glucose, 30 2,3-butanedione 2-monoxime; pH 7.3 when oxygenated at 37°C, ∼330 mOsm).

### Cell preparation

Left ventricular free wall was dissected (∼0.5 cm × 0.5 cm transmural fragments), and embedded in 4% low melting point agarose (Carl Roth, Karlsruhe, Germany). Slices of 300 µm thickness were obtained using a vibratome (model 7000-smz-2, Campden Instruments, Loughborough, UK), operated at a vibration frequency of 60 Hz, blade shift amplitude of 1.5 mm, and an advancement velocity of 50 µm/s, in oxygenated slicing solution at 4°C. Live tissue slices were stored in the fridge in solution B, and cardiomyocytes were isolated from slices on two days following organ excision, as described before.^27^ In addition to ventricular cardiomyocytes, cardiomyocytes were isolated from the left atrial free wall for comparisons, as described before.^27^

### Assessment of LTCC and NCX distribution

Cardiomyocytes were chemically fixed using 2% paraformaldehyde for 10 minutes at room temperature, then washed in phosphate-buffered saline (PBS), containing [in mM]: 137 NaCl, 2.7 KCl, 4.3 Na_2_HPO_4_, 1.47 KH_2_PO_4_; pH 7.4. Blocking and permeabilisation were performed for 1 h at room temperature with PBS containing 0.25% Triton X-100, 2.5% bovine serum albumin, and 5% goat serum (all Sigma-Aldrich, St. Louis, MO, USA). The cells were then incubated overnight at 4°C with primary antibodies (guinea pig anti-CaV1.2 [CACNA1C], dilution 1:200; Alomone Labs, Jerusalem, Israel; and mouse anti-NCX [C2C12], dilution 1:200; ThermoFisher, Waltham, MA, USA) in PBS containing 0.02% Triton-X, 1% bovine serum albumin, and 2.5% goat serum. Cells were then washed and incubated for 3 h at room temperature with secondary antibodies (goat anti-guinea pig IgG Alexa Fluor™ 488, dilution 1:500; and goat anti-mouse IgM Alexa Fluor™ 594, dilution 1:500; both ThermoFisher) in PBS containing 0.02% Triton-X, 1% bovine serum albumin, and 2.5% goat serum. Cells were subsequently shielded from light, washed with PBS, and mounted using Permafluor mounting medium (ThermoFisher).

Images of immuno-labelled cells were acquired using confocal laser scanning microscopy (Leica TCS SP8 X, Leica Microsystems, Wetzlar, Germany) with a HC PL APO CS2 63×/1.30 GLYC objective. For fluorescence excitation, appropriate “lines” (i.e. very narrow wavelength ranges) were picked from a white light laser. The same approach applied to imaging of Ca^2+^ dynamics inside TATS, see below. LTCC and NCX distribution were analysed using a custom Python script (available on request; to be made freely accessible upon acceptance of the manuscript), assessing the total number of photons in masks containing the sarcolemma of the entire cardiomyocyte, or one of two sub-domains: ‘surface membrane’ or ‘TATS’ only. For masks containing only the surface membrane, segmentation of the cell was performed using the NCX channel as a reference. An erosion of 3–4 pixels was then applied to isolate the cell’s inner region, which was subtracted from the whole-cell mask to obtain a surface membrane-specific mask. For the mask containing the TATS, segmentation of both LTCC and NCX channels was performed individually, using the Default/IJ_IsoData thresholding function in ImageJ/Fiji 2.9.0/1.53z. Since this automatic LTCC and NCX segmentation included signals from both the TATS and the surface membrane, the previously obtained surface membrane-specific mask was used to subtract the surface membrane area to obtain the TATS-only area.

### Ca^2+^ dynamics inside TATS

Assessment of [Ca^2+^]_TATS_ dynamics in individual, electrically stimulated ventricular cardiomyocytes was carried out at room temperature, using confocal imaging. Cells were placed in an RC-27NE2 recording chamber with field stimulation (Warner Instruments, Hamden, CT, USA), covered with a custom 3D-printed chamber lid to limit evaporation, and stimulated using a MyoPacer Field Stimulator (biphasic: 2 × 1.5 ms, ±12 V; IonOptix, Westwood, MA, USA).

To report extracellular Ca^2+^ levels, the non-membrane-permeable indicator Rhod-5N (tripotassium salt, 50 μM; AAT Bioquest, Sunnyvale, CA, USA) was used. The K*_D_* of Rhod-5N for Ca^2+^, assessed using recordings of the bath signal at increasing Ca^2+^ concentrations, was 0.4 mM (Supplementary Fig. S1). Recordings were performed in solution A, with Ca^2+^ concentration reduced to 0.6 mM (as a compromise between matching the indicator’s K*_D_* and maintaining an ionic environment as close to physiological as possible) and Mg^2+^ concentration reduced to 0.5 mM (as Rhod-5N also displays low sensitivity to Mg^2+^ ions).

For recordings in solution with increased viscosity (1.5 mPa·s, measured using MCR 303 Rheometer; Anton Paar, Graz, Austria), methylcellulose was added (final concentration 0.2%; Sigma-Aldrich). The impact of increased solution viscosity on apparent diffusion dynamics in TATS of isolated cells and tissue slices was assessed using fluorescence recovery after photobleaching (FRAP), as described before.^25^

Mechanical uncoupling was induced by addition of 40 μM para-amino-blebbistatin (Cayman Chemicals, Ann Arbor, MI, USA). In order to monitor the electrical activity of mechanically uncoupled cells, the cardiomyocytes were stained with 1 μM Di-4-ANEQ(F)PTEA (Potentiometric Probes, Farmington, CT, USA) as voltage sensor (Fig. 1A). A subset of experiments was conducted in the presence of 10 μM nifedipine (Sigma-Aldrich) as LTCC blocker. Rhod-5N was excited with a wavelength of 551 nm. The corresponding signal was recorded using a detection window of 558–607 nm; Di-4-ANEQ(F)PTEA was excited with a wavelength of 630 nm, and imaged using a detection window of 639–800 nm.

**Figure 1:**
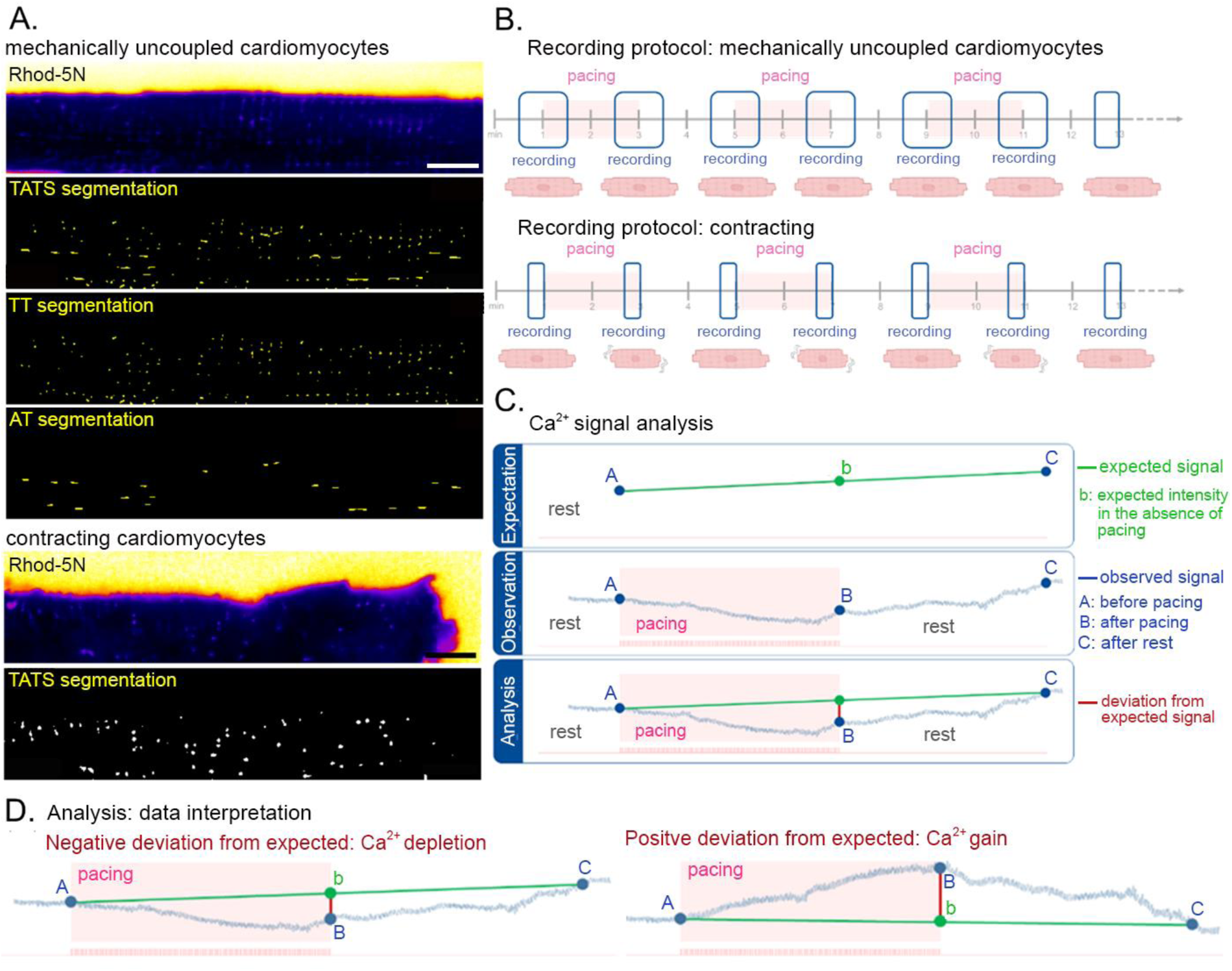
Experimental design and image analysis. **A:** Representative fields of view, showing parts of a cardiomyocyte in the presence of Rhod-5N as an extracellular Ca^2+^ indicator (sum projection), and segmentations of TATS, TT, and AT in mechanically uncoupled cardiomyocytes (top panels), and in freely contracting cardiomyocytes (bottom panels). Scale bars: 10 μm. **B:** Design of the experiment to probe relative changes in TATS Ca^2+^ levels ([Ca^2+^]_TATS_) during electrical pacing with increasing frequencies in mechanically uncoupled and contracting cardiomyocytes. Recordings were obtained in an interval mode to minimise phototoxic effects. Pale red background indicates periods of pacing. **C:** Calculation of the deviation from expected Ca^2+^ signal (explanation using an illustrative trace). This method of data extraction was implemented to correct for signal drift over time, due to environmental changes (incl. bleaching, evaporation, changes in temperature) or gradual changes in dye performance in biological system. **D:** Negative and positive deviation from expected Ca^2+^ signal and respective data interpretation.

Experiments on electrically stimulated, freely contracting cells were conducted in the presence of 0.425 mM of FITC-dextran (10 kDa; Sigma-Aldrich), serving as a bright fluorescent marker to facilitate automatic TATS detection during image analysis (Fig. 1A). The excitation wavelength was set to 488 nm, and the emission of FITC-dextran was collected in two adjacent spectral channels (498–516 nm and 521–559 nm) to obtain quasi-paired data with different noise observations for denoising. Addition of micromolar concentrations of dextran did not affect the measured viscosity of the solution.

For mechanically uncoupled cells, time-series were acquired with an acquisition rate of 36 frames/s (here a resonance scanner was used for confocal imaging). Recordings of freely-contracting cells were acquired using the standard galvo-scanner with an acquisition rate of 21 frames/s (slower frame rate when compared to mechanically uncoupled cell observations due to inclusion of additional channels). An interval imaging protocol was used (with periods of rest and stimulation at different pacing frequencies: 0.5, 1, 2, 3 Hz; Fig. 1B). The order of pacing frequencies was varied during individual experiments, and it was not found to affect results. In addition, a subset of time-matched control experiments was performed using the same imaging protocol, while no pacing was applied, to assess baseline Ca^2+^ dynamics at 0 Hz.

In addition to experiments using ventricular cardiomyocytes, a small subset of experiments was performed using atrial cardiomyocytes. These cells were recorded without supplementation with methylcellulose.

### Image analysis

For [Ca^2+^]_TATS_ analysis, changes in fluorescence intensity of the Ca^2+^ reporter Rhod-5N were quantified. All interval time-series from a single experiment were concatenated into a single time-series. TATS, TT, and AT segmentations were used to create masks which were then used to extract the signal from the time-series recordings. To generate segmentations of mechanically uncoupled cells, we calculated the sum-intensity projection of the Rhod-5N signal over time using Fiji 2.9.0/1.53t,^28^ and used Seg3D to annotate TATS, TT, and AT (Fig. 1A).^29^ Segmentation of TATS or TT did not include TT mouths (opening regions of the TT that connect to the extracellular space; defined as TT within 20 pixels from the cardiomyocyte surface (pixel size 0.18 µm^2^).

Recordings of cardiomyocytes that shifted in the X or Y plane were corrected by translation-only image registration using cross-correlation as the similarity metric, with each frame aligned to a mean-projection reference. Recordings of cardiomyocytes that showed signs of drift in the Z direction (altered focal plane) were discarded. To control for signal fluctuation due to evaporation, changes in temperature, and/or indicator signal drift over time, fluorescence intensities within TATS, TT, and AT masks were normalised to the signal from the bath, defined as the area >20 pixels from the surface membrane, where indicator and Ca^2+^ concentrations were assumed to remain constant. The mean bath fluorescence trace was fitted with a linear function, and the fluorescence signal in TATS, TT, and AT was normalised (divided) by the fitted bath values. Finally, the mean signal was calculated using 50 frames at different time points of the recordings (see Fig. 1B).

For the analysis of contracting cells, an automated TATS segmentation workflow was established that segmented each frame, rather than a single sum-intensity projection of the entire time-series. This was done in order to account for cell-movement-induced changes in the imaged sample. The whole-cell TATS signal was analysed, with no subdivision into TT and AT sources. First, we applied self-supervised Noise2Noise^30^ denoising using the implementation from the CSBDeep toolbox^31^ to the two acquired FITC-dextran signals. Then, we used a 2D UNet^32^ to segment TATS. To annotate ground truth data to train the network, we used the semi-automatic pixel classification workflow in ilastik^33^ to generate an initial segmentation, which we manually proofread and corrected in Seg3D. The network was trained using 3 manually segmented time-series of non-contracting cardiomyocytes and 1 manually segmented time-series of a contracting cardiomyocyte (using 50 frames). To obtain binary TATS segmentations, a threshold of 0.5 was applied (Fig. 1A). TATS Ca^2+^ fluorescence intensity was measured by averaging photon counts inside the TATS segmentation. The mean signal was calculated using 50 frames at different time points of the recordings.

Resting and pacing periods were annotated manually. Dynamics of the Ca^2+^ signal were analysed by extracting a value for ‘% deviation from expected Ca^2+^ signal’ at the end of each pacing and rest period. To calculate this, we fitted a linear regression from the time-point just before the start of each electrical pacing step to ∼120 seconds after the end of the pacing period, thus establishing a theoretically ‘expected’ signal trajectory reflecting a situation there was a continuous change in signal intensity (according to the null hypothesis that pacing has no effect on [Ca^2+^]_TATS_). The predicted intensity value, based on this linear fit (‘expected’ signal), at the end of any pacing run was then subtracted from the measured intensity value at the end of pacing, and the result was expressed as ‘% deviation from expected Ca^2+^ signal’ (Fig. 1C). This value was calculated by subtracting the expected from the observed values, and expressing it as a % of the expected value. A negative deviation from expected signals was taken to indicate TATS Ca^2+^ depletion, whereas a positive deviation was taken to indicate Ca^2+^ gain by TATS during pacing (Fig. 1D).

### In silico analyses

Electrophysiological effects of changes in [Ca^2+^]_TATS_ were studied in silico using the human ventricular cardiomyocyte model of Bartolucci et al.^34^ This model was selected because it reproduces the inverse relationship between action potential duration and [Ca^2+^]_o_. All simulations were performed with openCARP,^35^ employing an integration time step of 1 µs. Each simulation was run for 1,025 s. To exclude transient dynamics and study limit cycle characteristics, only the final 25 s of each simulation were used for analysis.

To investigate the effects of gradual Ca^2+^ depletion within the TATS, we introduced [Ca^2+^]_TATS_ explicitly in the model and considered a spatially heterogeneous distribution of LTCC and NCX. The TATS fraction of these channels was then exposed to [Ca^2+^]_TATS_, rather than bulk [Ca^2+^]_o_, as it would be in the original model formulation.

For the LTCC Ca^2+^ current (*I*_CaL_), all terms depending on [Ca^2+^]_o_ were duplicated. Consequently, the electrochemical driving force Ψ_Ca_ of Ca^2+^ across the surface membrane (SM), formulated using a Goldman-Hodgkin-Katz expression, was separated into a SM and a TATS component:

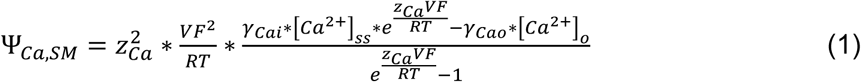

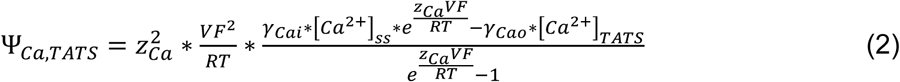

where z_Ca_ = +2 denotes the Ca^2+^ valence, V (mV) the transmembrane voltage, F (C*M^-1^) the Faraday constant, R (J*kM^-1^*K^-1^) the gas constant, T (K) the absolute temperature, γ_Cai_ = 1.2 and γ_Cao_ = 0.341 intracellular and extracellular activity coefficient for Ca^2+^, and [Ca^2+^]_ss_ (mM) the subsarcolemmal Ca^2+^ concentration.

The voltage-dependent (VD) and Ca^2+^-dependent (CD) components of *I*_CaL_ (*I*_CaL,VD_, *I*_CaL,VD,CaMK_, *I*_CaL,CD_, and *I*_CAL,CD,CaMK_) were calculated as weighted sums of Ca^2+^/calmodulin-dependent protein kinase (CaMK)-phosphorylated and non-phosphorylated channel populations, with weighting determined by the fraction of CaMK-phosphorylated channels ϕ_ICaL,CaMK_. Each component was split into SM and TATS contributions. This formulation preserves the original Bartolucci channel kinetics while enabling spatially heterogeneous weighting of LTCC between the SM and the TATS.

The parameter frac_ICaL,TATS_ denotes the fraction of LTCC localised in the TATS membrane, set to 0.75 unless stated otherwise (value based on data in Fig. 2). The total *I*_CaL_ was then expressed as the sum of these components:

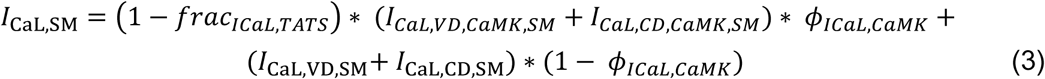

**Figure 2:**
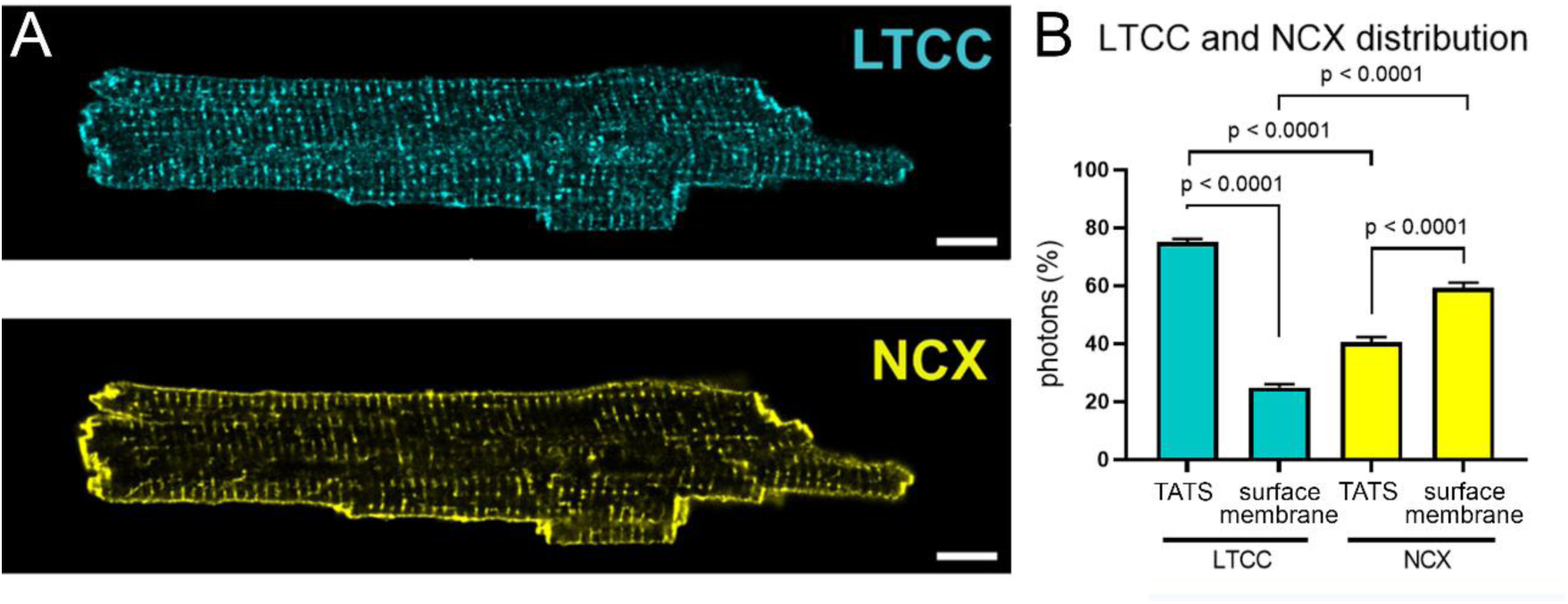
LTCC and NCX immunofluorescence in chemically fixed rabbit left ventricular cardiomyocytes. **A:** Representative confocal images of antibody-labelled LTCC (top) and NCX (bottom). **B:** Quantification of the distribution of photons collected from LTCC (cyan) and NCX (yellow) in TATS and surface membrane regions of the same cell. Data analysed using paired t-test; N = 35 cells / 3 hearts. Scale bars: 10 µm.

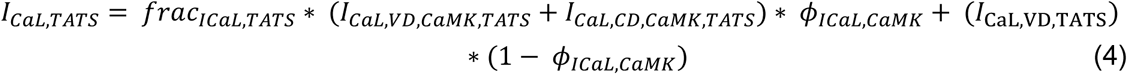

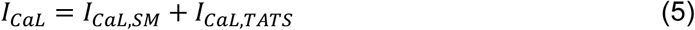

An analogous decomposition was applied to NCX. All NCX components directly or indirectly dependent on [Ca^2+^]_o_ were split into SM and TATS contributions. In particular, the Ca^2+^-binding rate to the intracellular regulatory Ca^2+^ site of NCX, namely k_1_, was reformulated as:

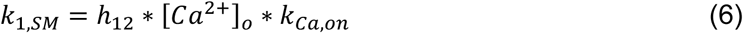

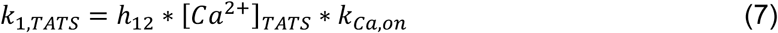

where h_12_ is a dimensionless coupling factor, and k_Ca,on_ = 1.5*10^6^ (mM*ms^-1^) the baseline association (on-rate) for binding to the intracellular regulatory site of NCX.

Consequently, the state variables x_2,Y_ – x_4,Y_, the exchanger states E_1,Y_ – E_4,Y_, fluxes J_NaCa,Na,Y_ and J_NaCa,Ca,Y_, and the resulting exchanger current *I*_NaCa,Y_ were all split into SM and TATS components. Here, the index Y denotes either the bulk intracellular (i) or sub-sarcolemmal (ss) Ca^2+^ compartments, which are coupled via a diffusive Ca^2+^ flux J_diff_. The newly introduced parameter frac_INaCa,TATS_ denotes the fraction of NCX localised in the TATS membrane, set to 0.4 (value based on data in Fig. 2). All remaining equations were adopted from the model of Bartolucci et al.^34^ and the parental O’Hara et al. model.^36^

A reduction of [Ca^2+^]_TATS_ from 1.8 mM to 1.6 mM was simulated to investigate the effects of Ca^2+^ depletion in the TATS on the action potential and intracellular Ca^2+^ transient. This degree of depletion is based on experimental data obtained at a pacing frequency of 3 Hz, where the observed Ca^2+^ depletion in TATS was ∼10 %, as calculated based on Rhod-5N calibration curve (Supplementary Fig. S1). Control traces were obtained with [Ca^2+^]_TATS_ of 1.8 mM. Finally, the human active tension model of Land et al.^37^ was coupled with the Bartolucci et al. cardiomyocyte electrophysiology model to predict consequences of TATS Ca^2+^ depletion on active tension development.

### Data analysis

Data are presented as mean ± SEM, and analyses were performed using Prism 10 (GraphPad, San Diego, CA, USA). Data were analysed using paired t-tests or one-way ANOVA with linear trend analyses; as indicated in figure legends. A *p*-value < 0.05 was taken to indicate a statistically significant difference between means, or to assess the presence of a trend.

## RESULTS

### LTCC and NCX distribution in rabbit left ventricular cardiomyocytes

LTCC and NCX are the primary pathways for transmembrane Ca²⁺ influx and extrusion, respectively. LTCC and NCX distribution in TATS and surface membrane was analysed using immunohistochemical labelling. Relative LTCC abundance, calculated as the % of photons collected from either membrane subsystem, was higher in TATS than in surface membrane in confocal image sections (74.37±1.02% in TATS vs 25.63±1.02% in surface membrane, N = 35 cells / 3 hearts, p < 0.0001), while NCX was more abundant in the surface membrane compared to TATS (40.59±1.74% in TATS vs 59.41±1.74% in surface membrane, N = 35 cells / 3 hearts, p < 0.0001). When analysing each of these sub-domains separately, the TATS mask contained a higher proportion of LTCC-associated photons compared to NCX (p<0.0001), whereas the surface membrane mask contained a higher proportion of NCX-associated photons compared to LTCC (p<0.0001, Fig. 2).

### Pacing rate dependent effects on intraluminal TATS Ca^2+^ dynamics in mechanically uncoupled rabbit left ventricular cardiomyocytes

To assess [Ca^2+^]_TATS_ dynamics at different pacing frequencies while excluding effects of cell contractions, changes in fluorescence intensity of the extracellular Ca^2+^ indicator Rhod-5N were monitored in TATS of mechanically uncoupled cardiomyocytes. Experiments were conducted in a solution A with [Ca^2+^] reduced to 0.6 mM and [Mg^2+^] reduced to 0.5 mM, supplemented with 0.2% w/v methylcellulose to increase viscosity, to better mimic the diffusive environment that cells would be exposed to in tissue (Supplementary Fig. S2). In particular, when measuring apparent diffusion speeds in TATS using fluorophore-conjugated dextran particles and FRAP microscopy,^24,25^ increasing solution viscosity from 1 mPa·s to 1.5 mPa·s (after supplementation with methylcellulose) gives rise to FRAP time constants (*τ*) that are no longer significantly different from values seen in tissue slices (cell at 1 mPa·s: 0.48±0.02 s, cell at 1.5 mPa·s: 0.66±0.03 s, slice: 0.8±0.13 s, N = 51–65 cells / 8 slices / 3 hearts, p = 0.002 for cell at 1 mPa·s vs 1.5 mPa·s, p = 0.002 for cell at 1 mPa·s vs slice, p = 0.43 for cell at 1.5 mPa·s vs slice; Supplementary Fig. S2).

We analysed the Ca^2+^ dynamics in whole-cell TATS, as well as separately in TT and AT. The [Ca^2+^]_TATS_ dynamics were analysed as ‘% deviation from expected Ca^2+^ signal’ at the end of each pacing period (see Fig. 1). In mechanically uncoupled ventricular cardiomyocytes, electrical pacing was associated with an apparent Ca^2+^ depletion in TATS (0 Hz: 0.61±0.5%, 0.5 Hz: 0.94±0.5%, 1 Hz: 0.25±0.4%, 2 Hz: -1.31±0.8%, 3 Hz: -2.96±2.3%, N = 4–51 cells/ 6 hearts, p = 0.033), specifically in TT (0 Hz: 0.51±0.5%, 0.5 Hz: 0.75±0.5%, 1 Hz: 0.08±0.6%, 2 Hz: -1.87±0.6%, 3 Hz: -2.9±1.4%, N = 5–51 cells / 6 hearts, p = 0.027), with no significant effect of pacing on the highly variable AT luminal Ca^2+^ dynamics (0 Hz: 0.11±0.9%, 0.5 Hz: - 2.22±1.2%, 1 Hz: 0.76±1.2%, 2 Hz: -0.43±2.3%, 3 Hz: -7.49±3.8%, N = 3–33 cells / 6 hearts, p = 0.53). The pacing-induced reduction in [Ca^2+^]_TATS_, seen in TATS and TT, scaled with increasing pacing frequency (Fig. 3).

**Figure 3:**
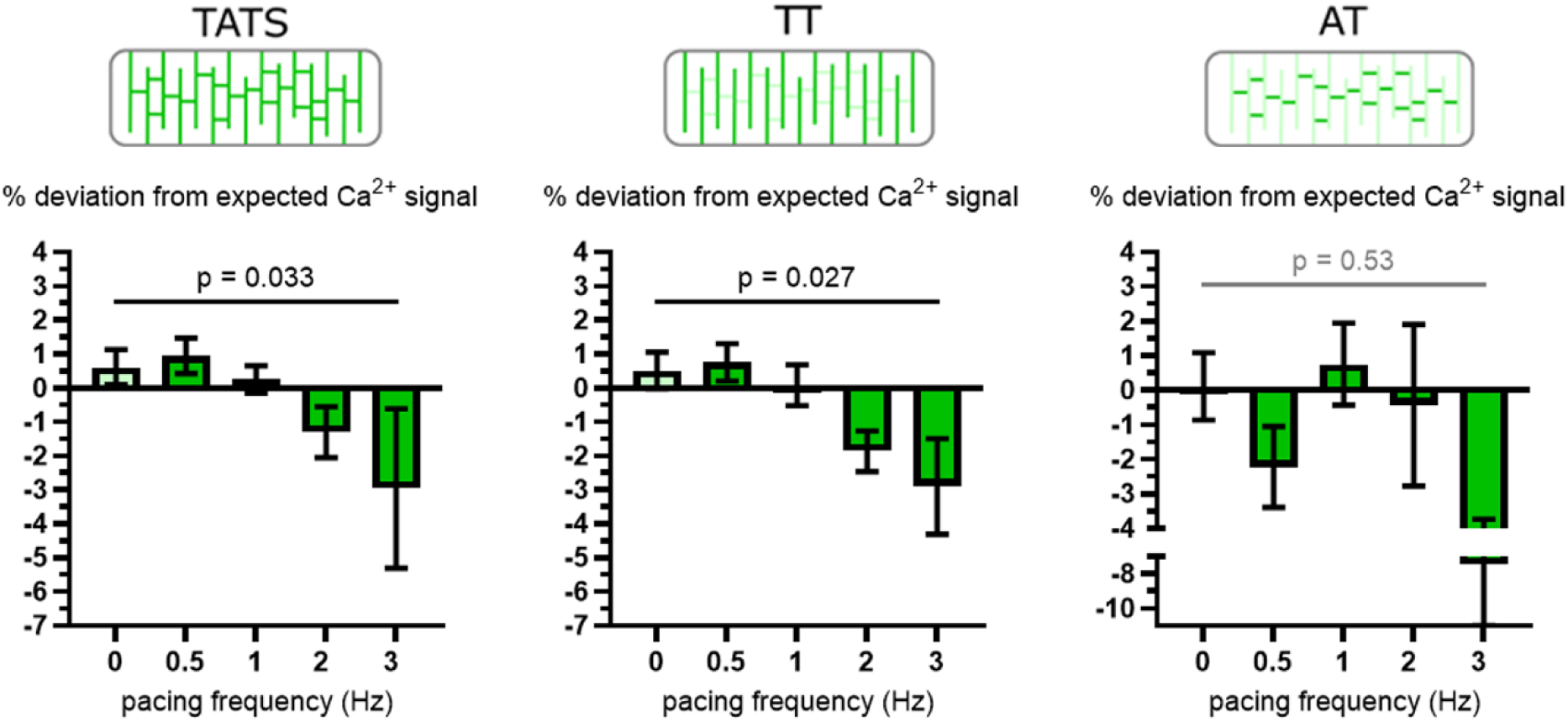
Electrical pacing rate and Ca^2+^ dynamics in TATS, TT, and AT of mechanically uncoupled rabbit left ventricular cardiomyocytes. Data for ‘0 Hz’ were obtained in a separate experiment aimed as assessing baseline Ca^2+^ dynamics in unstimulated cardiomyocytes, using the same experimental design and image analysis pipelines as used for data shown for 0.5–3 Hz. Data analysed using one-way ANOVA with a test for linear trend; N = 3–51 cells / 6 hearts.

To determine whether Ca^2+^ depletion in TATS and TT requires transmembrane fluxes through LTCC, nifedipine was administered in separate experiments to block LTCC activity. In the presence of nifedipine, there was no significant effect of pacing on intraluminal Ca^2+^ dynamics in TATS (0 Hz: 0.51±0.5%, 0.5 Hz: 0.86±0.8%, 1 Hz: -0.92±0.7%, 2 Hz: 2.17±0.8%, 3 Hz: 1.65±1.3%, N = 11–50 cells / 9 hearts, p = 0.74), TT (0 Hz: 0.51±0.5%, 0.5 Hz: 0.75±0.5%, 1 Hz: 0.08±0.6%, 2 Hz: -1.87±0.6%, 3 Hz: -2.9±1.4%, N = 5–51 cells / 9 hearts, p = 0.24), or AT (0 Hz: 0.11±0.9%, 0.5 Hz: 0.71±1.7%, 1 Hz: -1.85±2%, 2 Hz: 1.47±1.6%, 3 Hz: -0.97±4.5%, N = 10–33 cells / 9 hearts, p = 0.93) of mechanically uncoupled cells (Fig. 4).

**Figure 4:**
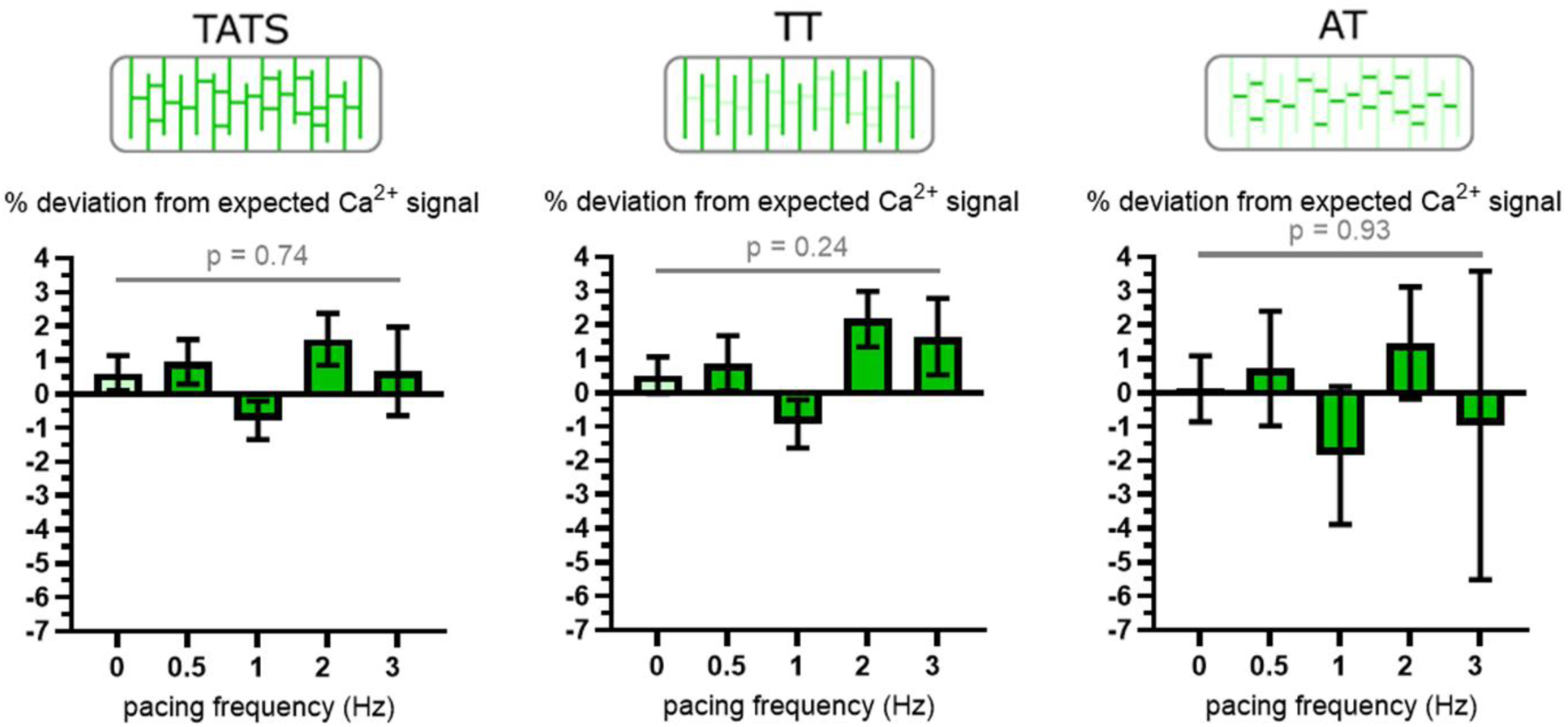
Electrical pacing rate and Ca^2+^ dynamics in TATS, TT, and AT of mechanically uncoupled rabbit left ventricular cardiomyocytes with pharmacological LTCC inhibition. Cells were imaged in the presence of nifedipine (10 µM). Data for ‘0 Hz’ were obtained in a separate experiment aimed as assessing baseline Ca^2+^ dynamics in unstimulated cardiomyocytes, using the same experimental design and image analysis pipelines as used for data shown for 0.5–3 Hz. Data analysed using one-way ANOVA with a test for linear trend; N = 10–50 cells / 9 hearts.

In addition to experiments on ventricular cardiomyocytes, we also assessed [Ca^2+^]_TATS_ in atrial cardiomyocytes. Similar to ventricular cells, analysis of immunohistochemical labelling of fixed cells revealed that LTCC is more abundant in TATS, compared to surface membrane regions (67.6±0.78% in TATS vs 32.4±0.78% in surface membrane, N = 71 cells / 3 hearts, p < 0.001), with no significant difference in the distribution of NCX between the two membrane sub-domains (50.48±1.03% in TATS vs 49.52±0.78% in surface membrane, N = 71 cells / 3 hearts, p > 0.99; Supplementary Fig. S3). When analysing each sub-domain separately, the TATS mask contained a higher proportion of LTCC-associated photons compared to NCX (p<0.001), whereas the surface membrane mask contained a higher proportion of NCX-associated photons compared to LTCC (p<0.001), as seen in ventricular cells.

Electrical pacing of mechanically uncoupled atrial cardiomyocytes led to a pacing rate dependent Ca^2+^ depletion in TATS, as seen in ventricular cells. This depletion was significant for TATS (0.5 Hz: 1.87±1.6%, 1 Hz: -0.13±1.63%, 2 Hz: -2.93±2.3%, N = 32–33 cells / 11 hearts, p = 0.039), TT (0.5 Hz: 3.02±2%, 1 Hz: 1.59±1.2%, 2 Hz: -3.3±1.8%, N = 32-33 cells / 11 hearts, p = 0.013), and AT (0.5 Hz: 1.91±1%, 1 Hz: 0.64±1.4%, 2 Hz: -2.82±1.4%, N = 32–33 cells / 11 hearts, p = 0.01), suggesting that the effect is independent of cardiac chambers (Supplementary Fig. S4).

### Lack of pacing-rate dependent effects on intraluminal TATS Ca^2+^ dynamics in contracting rabbit left ventricular cardiomyocytes

In order to assess the impact of cardiomyocyte contractions on [Ca^2+^]_TATS_ dynamics, cells were allowed to contract freely during pacing, and Ca^2+^ reporter fluorescence was analysed as described above. Experiments were conducted at room temperature, and contractions were reliably inducible at pacing rates up to 2 Hz. Cells contracted with a shortening amplitude of 5–15 % (assessed using TT-TT distance changes as a surrogate measure of sarcomere shortening). The entire TATS was analysed, without distinction between TT and AT. In contrast to observations in mechanically uncoupled cells, we observed no evidence of TATS Ca^2+^ depletion in contracting cells (0 Hz: 0.61±0.5%, 0.5 Hz: -2.21±1%, 1 Hz: 1.95±1.2%, 2 Hz: - 1.74±2.1%, N = 6–56 cells / 10 hearts, p = 0.51; Fig. 5).

**Figure 5:**
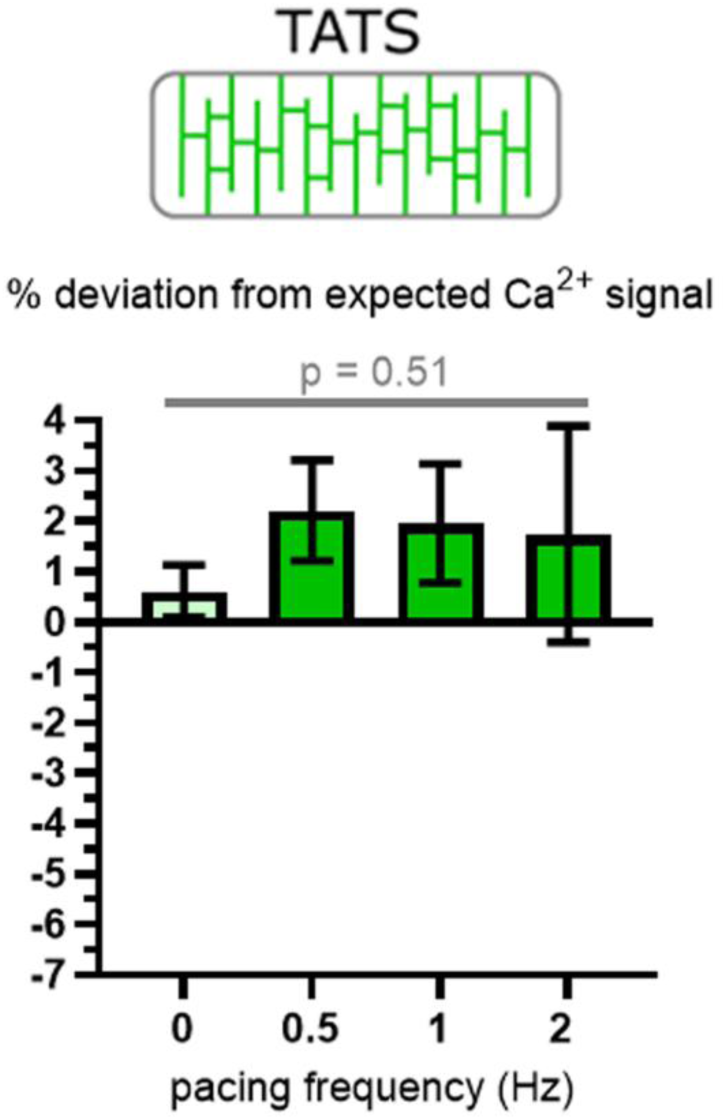
Electrical pacing rate and Ca^2+^ dynamics in TATS of electrically stimulated, freely contracting rabbit left ventricular cardiomyocytes. Data for ‘0 Hz’ were obtained in a separate experiment aimed as assessing baseline Ca^2+^ dynamics in unstimulated cardiomyocytes, using the same experimental design and image analysis pipelines as used for data shown for 0.5–3 Hz in Fig. 3 and 4. Data analysed using one-way ANOVA with a test for linear trend; N = 6–56 cells/ 10 hearts.

### Computational modelling of the effects of Ca^2+^ depletion in TATS on cardiomyocyte function

Computational modelling, incorporating the relative LTCC and NCX distribution in TATS and surface membrane (data in Fig. 2) revealed that a decrease in [Ca^2+^]_TATS_ to the level seen in mechanically uncoupled cardiomyocytes (based on the calibration curve of the dye, we estimate the decrease in [Ca^2+^]_TATS_ to be ∼10% at 3 Hz) leads to a decrease in intracellular Ca^2+^ transient amplitude, and a decrease in active tension development, with no significant effects on the action potential amplitude or duration (Fig. 6).

**Figure 6:**
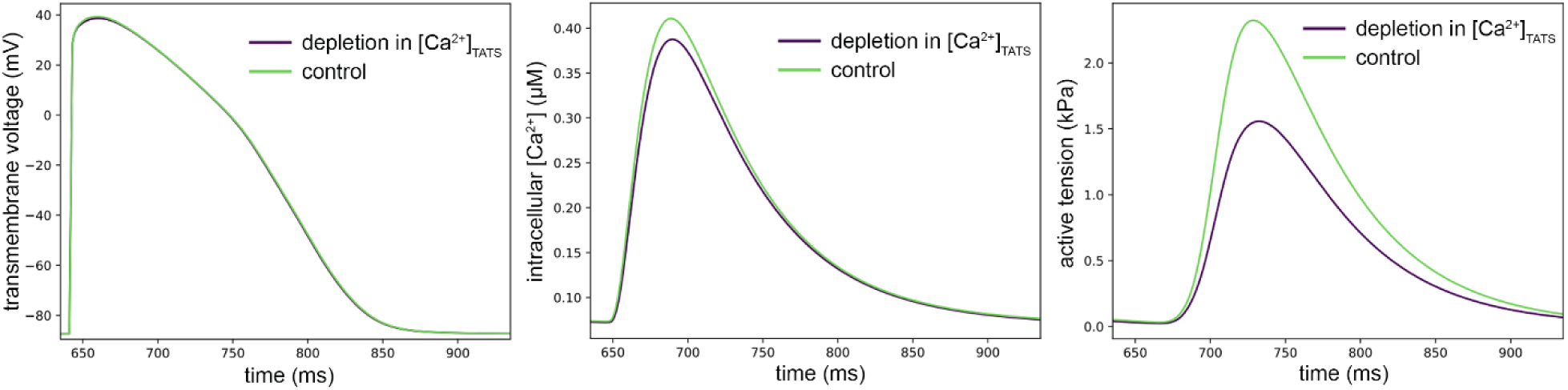
Computational simulation of the effects of TATS Ca^2+^ depletion on cardiomyocyte function in mechanically uncoupled cardiomyocytes. A decrease in [Ca^2+^]_TATS_ from 1.8 mM (control) to 1.6 mM (mimicking 3 Hz pacing in mechanically uncoupled cells) does not significantly affect action potential properties, but leads to a decrease in intracellular Ca^2+^ transient amplitude and a decrease in active tension development. Ca^2+^ concentration in bulk extracellular space was maintained at 1.8 mM.

## DISCUSSION

Exploring extracellular Ca^2+^ reporter fluorescence changes in TATS of rabbit ventricular cardiomyocytes, we provide evidence that both electrical pacing and mechanical activity of cardiomyocytes affect [Ca^2+^]_TATS_ dynamics. We show that pacing at increasing frequencies leads to apparent Ca^2+^ depletion in TATS of mechanically uncoupled cardiomyocytes (in ventricular cells specifically in TT; in atrial cardiomyocytes both in TT and AT). We attribute these observations to an uneven distribution of LTCC and NCX across the surface membrane of cardiomyocytes, with a substantially higher abundance of LTCC, but not of NCX, in TATS, compared to the outer surface membrane. This can give rise to a net redistribution of Ca^2+^ from TATS lumen to bulk extracellular space – a suggestion supported by the evidence that blocking LTCC abolishes the pacing-induced Ca^2+^ depletion in TATS of non-contracting cells. Our *in silico* analyses reveal that – if unrectified – the levels of Ca^2+^ depletion seen experimentally in TATS at physiological (for rabbit) pacing rates would decrease intracellular Ca^2+^ transient amplitudes and active tension development. This TATS Ca^2+^ depletion appears to be countered by cell contraction-induced mechanical advection of TATS content, as it is not seen in electrically stimulated, freely shortening cardiomyocytes.

While no experimental evidence for Ca^2+^ depletion in TATS has been published prior to this study, earlier work using Ca^2+^–sensitive microelectrodes and colorimetric assays has provided indication that near-membrane Ca^2+^ depletion may occur as a consequence of cardiomyocyte electrical activity in multicellular preparations.^16,18,19,21^ The degree of Ca^2+^ depletion seen in these previous experimental studies ranged from 0.2–10%, and was up to 19.8% in computational work.^22,23^ Here, based on the dye calibration curve, we estimate the decrease in [Ca^2+^]_TATS_ to be ∼10% at 3 Hz, which is in keeping with previous estimates. This degree of Ca^2+^ depletion in TATS would be sufficient to negatively impair excitation–contraction coupling, as seen in our computational modelling.

We were able to dissect Ca^2+^ dynamics in TT vs AT in electro-mechanically uncoupled ventricular and atrial cardiomyocytes. Classically, AT are not thought to participate in excitation−contraction coupling in ventricular cells, and we did not observe a statistically significant impact of electrical pacing on Ca^2+^ levels in AT. However, AT in atrial cells are major Ca^2+^ release hubs.^2^ This is in keeping with the observed pacing-induced Ca^2+^ depletion in AT of atrial cardiomyocytes, and should be followed up in more detail in future work.

Our data suggest that the development of [Ca^2+^]_TATS_ gradients is countered in contracting cells by TATS content agitation and advection. While advection-assisted diffusion, driven by TT deformation during cell contraction and stretch (and AT deformation during stretch), has been reported,^24,25^ the possibility that cell contractions can affect intraluminal [Ca^2+^]_TATS_ dynamics had not been previously substantiated. Pathological remodelling, especially if combined with reduced contractile activity, may limit the extent to which TATS deformation can mitigate pacing rate dependent Ca^2+^ depletion. This will be subject to further research, also in view of the frequently encountered structural remodelling of the TATS, whose relevance of beating-rate dependent TATS [Ca^2+^] dynamics remain to be elucidated.

Our study has a number of limitations. Measurements have been performed at room temperature, and it is likely that diffusive dynamics we observed were slower than at body temperature. Furthermore, we were not able to achieve pacing rates higher than 2–3 Hz, and we could not systematically apply preloads to isolated cells – so the full extent of pacing-induced and advection-curbed changes in TATS content exchange is likely to be larger. Also, our experiments were performed at lower than physiological Ca^2+^ concentrations (0.6 mM). This was done to match the K*_D_* of available indicators. In contracting cells, we did not differentiate between AT and TT, and quantified TATS Ca^2+^ dynamics irrespective of location, an approach dictated by the need to avoid motion-induced artefacts in our analyses. Our studies have been performed using isolated cardiomyocytes. While constituting a well-established model with many benefits, enzymatic isolation interferes with the extracellular matrix lining cell membranes, which can disrupt the ability of the basement membrane to buffer Ca^2+^.^21^ Future studies would ideally explore multicellular preparations in more detail. In any case, we did mimic the apparently higher viscosity of solutions in tissue micro-domains using methylcellulose to match interstitial fluid properties in live tissue slices. Finally, in future studies the dynamics of Ca^2+^ depletion in TATS should be investigated over longer time-scales, to assess whether a new steady state is eventually reached, and if so, to explore experimentally how this affects intracellular Ca transients and sarcoplasmic reticulum storage.

In summary, we provide experimental evidence for a pacing rate-dependent depletion of Ca^2+^ inside TATS, which is countered by mechanically induced advection of TATS luminal content in contracting cells. This interplay of electrical and mechanical effects supports [Ca^2+^]_TATS_ homeostasis. Whether or not this would be disturbed during pathological remodelling of TATS structure and cardiomyocyte function remains to be elucidated.

## Acknowledgements

We would like to thank Stefanie Perez-Feliz, Cinthia Walz, Pia Iaconniani, Manuel Koch, and Jonas Heer for technical assistance with cell isolation, and Alina Semenjakin and Phil Henneken for assistance with imaging protocol optimisation. We acknowledge the SCI-MED imaging facility at the Institute for Experimental Cardiovascular Medicine in Freiburg for access to the confocal microscope. This research was supported by the German Research Foundation Emmy Noether programme (396913060 to EARZ). JG, JM, CZJ, PK, and EARZ are members of the Collaborative Research Centre SFB1425 of the German Research Foundation (422681845).

## Supplementary figures

**Figure S1:**
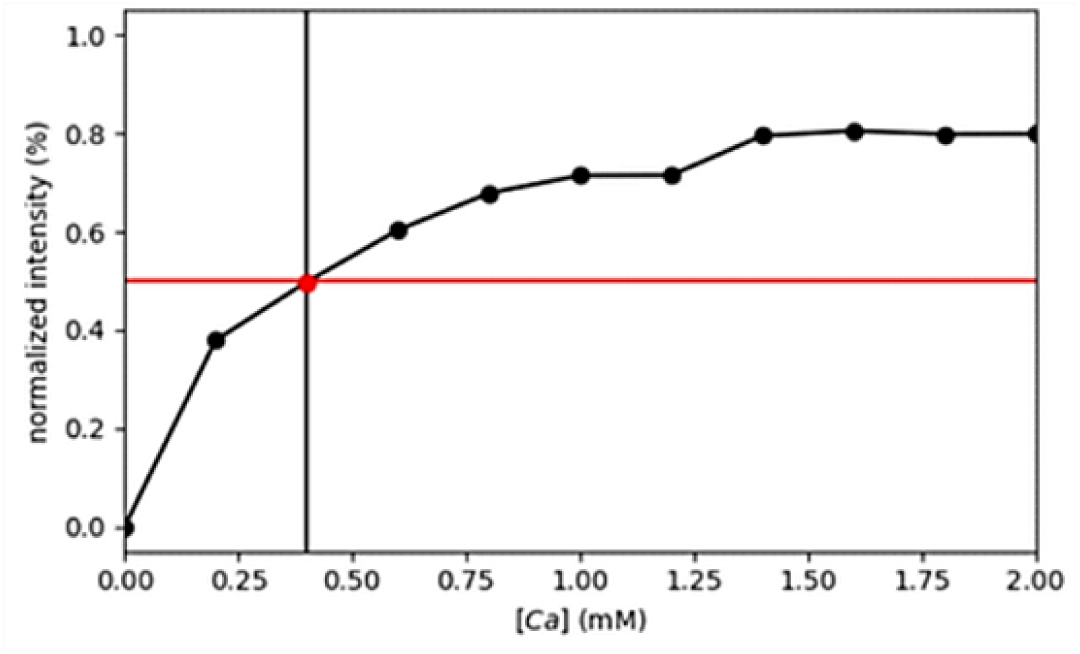
K_D_ measurements of Rhod-5N in solution supplemented with methylcellulose (0.2% w.v). K_D_ = 0.40 mM.

**Figure S2:**
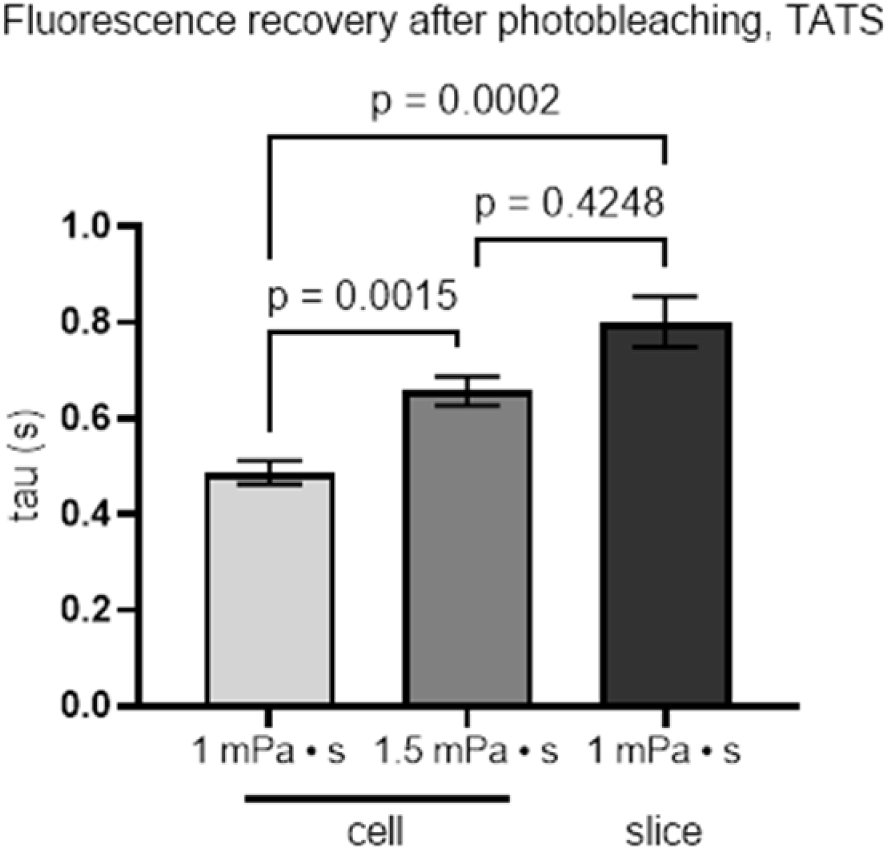
Ca^2+^ indicator (Rhod-5N)-based assessment of apparent diffusion speed in TATS, observed by FRAP (as described before^24^), in isolated cardiomyocytes and tissue slices. At a viscosity of 1 mPa·s, the time constant (τ) is substantially lower in single cells (left bar) compared to tissue (right bar). Elevating viscosity to 1.5 mPa·s (buffer supplemented with 0.2% methylcellulose) raises τ in single cells (middle bar) to levels no longer significantly different from tissue. Data analysed using Kruskal-Wallis test with Dunn’s post-hoc test; N = 51–65 cells / 8 slices / 3 hearts.

**Figure S3:**
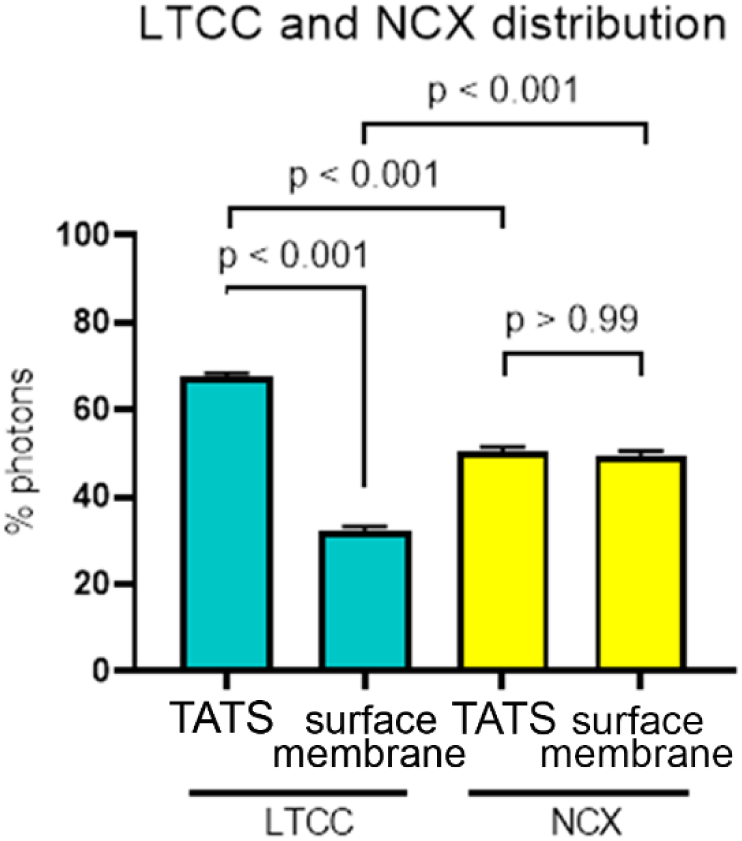
LTCC and NCX immunofluorescence in chemically fixed rabbit left atrial cardiomyocytes. Quantification of the distribution of photons, collected from LTCC (cyan) and NCX (yellow) in TATS and surface membrane regions of the same cell. Data analysed using paired t-test; N = 71 cells / 3 hearts

**Figure S4:**
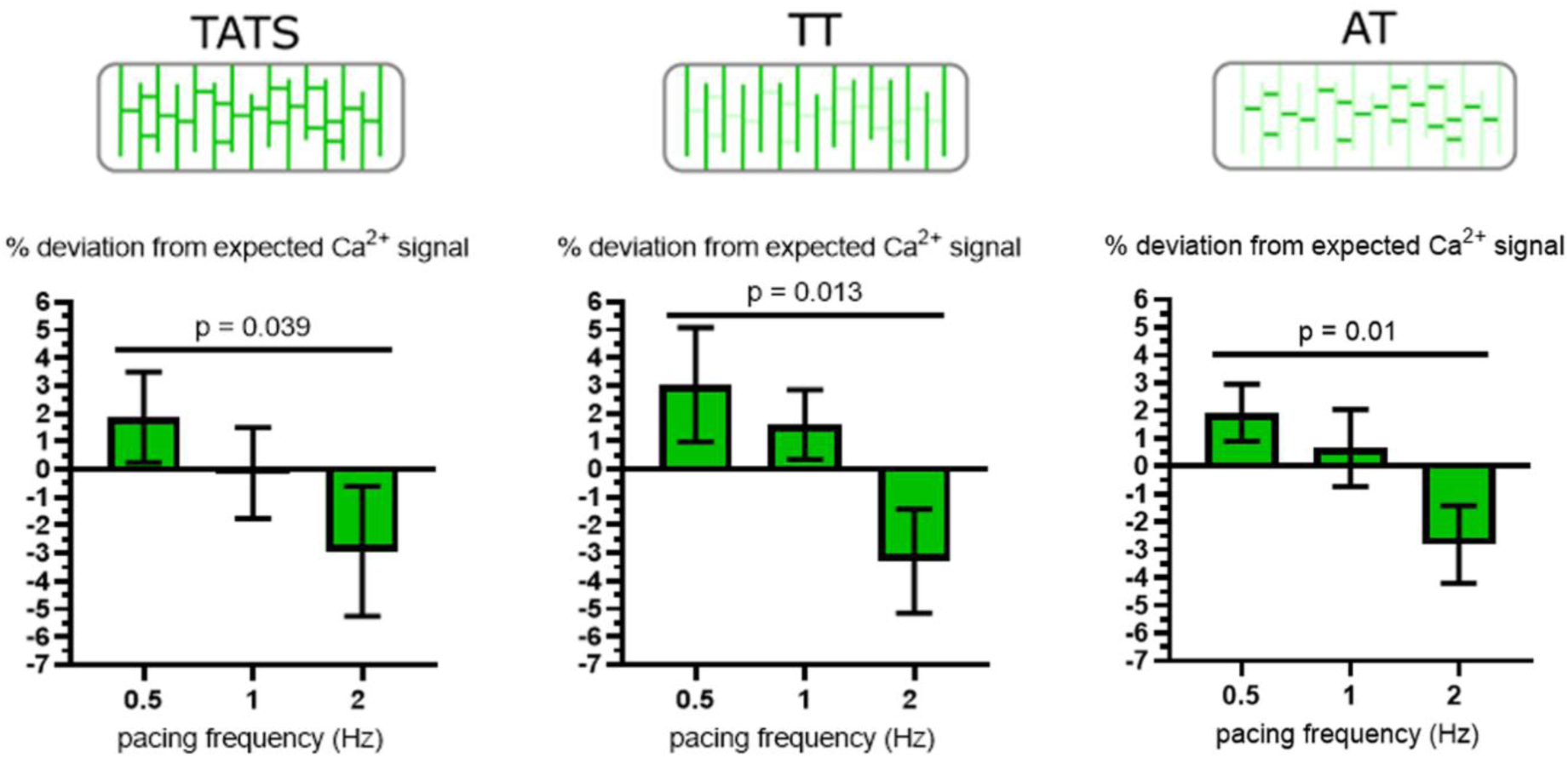
Electrical pacing rate and Ca^2+^ dynamics in TATS, TT, and AT of mechanically uncoupled rabbit left atrial cardiomyocytes. Data were analysed using one-way ANOVA with a test for linear trend; N = 32–33 cells / 11 hearts.

## REFERENCES

1. Brette F, Orchard C. T-tubule function in mammalian cardiac myocytes. Circ Res 2003;92(11):1182–1192

2. Brandenburg S, Kohl T, Williams GSB, Gusev K, Wagner E, Rog-Zielinska EA, Hebisch E, Dura M, Didie M, Gotthardt M, Nikolaev VO, Hasenfuss G, Kohl P, Ward CW, Lederer WJ, Lehnart SE. Axial tubule junctions control rapid calcium signaling in atria. J Clin Invest 2016;126(10):3999–4015

3. Setterberg IE, Le C, Frisk M, Li J, Louch WE. The physiology and pathophysiology of t-tubules in the heart. Front Physiol 2021;12:1–21

4. Scardigli M, Crocini C, Ferrantini C, Gabbrielli T, Silvestri L, Coppini R, Tesi C, Rog-Zielinska EA, Kohl P, Cerbai E, Poggesi C, Pavone FS, Sacconi L. Quantitative assessment of passive electrical properties of the cardiac T-tubular system by FRAP microscopy. Proc Natl Acad Sci USA 2017;114(22):5737–5742

5. Crossman DJ, Ruygrok PR, Soeller C, Cannell MB. Changes in the organization of excitation-contraction coupling structures in failing human heart. PLoS One 2011;6(3)

6. Bers DM. Cardiac excitation–contraction coupling. Nature 2002;415(6868):198–205

7. Orchard CH, Pásek M, Brette F. The role of mammalian cardiac T-tubules in excitation-contraction coupling: experimental and computational approaches. Exp Physiol 2009;94(5):509–519

8. Scriven DRL, Dan P, Moore EDW. Distribution of proteins implicated in excitation-contraction coupling in rat ventricular myocytes. Biophys J 2000;79(5):2682–2691

9. Pásek M, Šimurda J, Bébarová M, Christé G. Divergent estimates of the ratio between Na^+^-Ca^2+^ current densities in t-tubular and surface membranes of rat ventricular cardiomyocytes. J Cell Sci 2021;134(14):1–7

10. Vermij SH, Abriel H, Kucera JP. A fundamental evaluation of the electrical properties and function of cardiac transverse tubules. Biochim Biophys Acta - Mol Cell Res 2020;1867(3):118502

11. Kong CHT, Rog-Zielinska EA, Kohl P, Orchard CH, Cannell MB. Solute movement in the T-tubule system of rabbit and mouse cardiomyocytes. Proc Natl Acad Sci USA 2018;115(30):7073–7080

12. Rog-Zielinska EA, Moss R, Kaltenbacher W, Greiner J, Verkade P, Seemann G, Kohl P, Rog-Zielinska EA. Nano-scale morphology of cardiomyocyte t-tubule/sarcoplasmic reticulum junctions revealed by ultra-rapid high-pressure freezing and electron tomography. J Mol Cell Cardiol 2021;153:86–92

13. Shepherd N, McDonough HB. Ionic diffusion in transverse tubules of cardiac ventricular myocytes. Am J Physiol - Heart Circ Physiol 1998;275:852–860

14. Kubasov I V., Stepanov A, Bobkov D, Radwanski PB, Terpilowski MA, Dobretsov M, Gyorke S. Sub-cellular electrical heterogeneity revealed by loose patch recording reflects differential localization of sarcolemmal ion channels in intact rat hearts. Front Physiol 2018;9:1–9

15. Scriven DR, Klimek A, Lee KL ME. The molecular architecture of calcium Microdomains in rat cardiomyocytes. Ann N Y Acad Sci 2002;976(1):488–499

16. Bers DM, MacLeod KT. Cumulative depletions of extracellular calcium in rabbit ventricular muscle monitored with calcium-selective microelectrodes. Circ Res 1986;58(6):769–782

17. Bers DM, Peskoff A. Diffusion around a cardiac calcium channel and the role of surface bound calcium. Biophys J 1991;59(3):703–721

18. Hilgemann DW. Extracellular calcium transients at single excitations in rabbit atrium measured with tetramethylmurexide. J Gen Physiol 1986;87(5):707–735

19. Hilgemann DW, Langer GA. Transsarcolemmal calcium movements in arterially perfused rabbit right ventricle measured with extracellular calcium-sensitive dyes. Circ Res 1984;54(4):461–467

20. Shattock MJ and Bers DM. Rat vs rabbit ventricle: Ca^2+^ flux and intracellular Na^+^ assessed by ion-selective microelectrodes. Am J Physiol 1989;256(25):813–822

21. Bers D. Early transient depletion of extracellular Ca^2+^ during individual cardiac muscle contractions. Am J Physiol 1983;244(3):462–868

22. Pásek M, Šimurda J, Christé G, Orchard CH. Modelling the cardiac transverse-axial tubular system. Prog Biophys Mol Biol 2008;96(1–3):226–243

23. Pásek M, Šimurda J, Orchard CH, Christé G. A model of the guinea-pig ventricular cardiac myocyte incorporating a transverse–axial tubular system. Prog Biophys Mol Biol 2008;96(1–3):258–280

24. Rog-Zielinska EA, Scardigli M, Peyronnet R, Zgierski-Johnston CM, Greiner J, Madl J, O’Toole E, Morphew M, Hoenger A, Sacconi L, Kohl P. Beat-by-beat cardiomyocyte T-tubule deformation drives tubular content exchange. Circ Res 2021;128(2):203–215

25. Greiner J, Dente M, Orós-Rodrigo S, Cameron BA, Madl J, Kaltenbacher W, Kok T, Zgierski-Johnston CM, Peyronnet R< Kohl P, Sacconi L, Rog-Zielinska EA. Different effects of cardiomyocyte contractile activity on transverse and axial tubular system luminal content dynamics. J Mol Cell Cardiol 2024;197:125–135

26. McNary TG, Spitzer KW, Holloway H, Bridge JHB, Kohl P, Sachse FB. Mechanical modulation of the transverse tubular system of ventricular cardiomyocytes. Prog Biophys Mol Biol 2012;110(2–3):218–225

27. Greiner J, Schiatti T, Kaltenbacher W, Dente M, Semenjakin A, Kok T, Fiegle DJ, Seidel T, Ravens U, Kohl P, Peyronnet R, Rog-Zielinska EA. Consecutive-day ventricular and atrial cardiomyocyte isolations from the same heart: shifting the cost–benefit balance of cardiac primary cell research. Cells 2022;11(2):233

28. Schindelin J, Arganda-Carreras I, Frise E, Kaynig V, Longair M, Pietzsch T, Preibisch S, Rueden C, Saalfeld S, Schmid B, inevez J-Y, While DJ, Hartenstein V, Eliceiri K, Tomancak P, Cardona A. Fiji: An open-source platform for biological-image analysis. Nat Methods 2012;9(7):676–82.

29. Seg3D: Scientific Computing and Imaging Institute, University of Utah. [Software]. Version 2.2.1. Available from: https://www.seg3d.org/.

30. Lehtinen J, Munkberg J, Hasselgren J, Laine S, Karras T, Aittala M, Aila T. Noise2Noise: Learning image restoration without clean data. 35th Int Conf Mach Learn 2018;7(3):4620–4631

31. Weigert M, Schmidt U, Boothe T, Müller A, Dibrov A, Jain A, Wilhelm B, Schmidt D, Broaddus C, Culley S, Rocha-Martins M, Segovia-Miranda F, Norden C, Henriques R, Zerial M, Solimena M, Rink J, Tomancak P, Royer L, Jug F, Myers EW. Content-aware image restoration: pushing the limits of fluorescence microscopy. Nat Methods 2018;15(12):1090–1097

32. Ronneberger O, Fischer P BT. U-Net: Convolutional Networks for Biomedical Image Segmentation. In: Navab N, Hornegger J, Wells W, Frangi A. Medical image computing and computer-assisted intervention – MICCAI 2015. Lecture Notes in Computer Science 2015; 9351. Cham. Springer.

33. Berg S, Kutra D, Kroeger T, Straehle CN, Kausler BX, Haubold C, Schiegg M, Ales J, Beier T, Rudy M, Eren K, Cervantes JI, Xu B, Beuttenmueller F, Wolny A, Zhang C, Koethe U, Hamprecht FA, Kreshuk A. Ilastik: Interactive machine learning for (bio)image analysis. Nat Methods 2019;16(12):1226–1232

34. Bartolucci C, Passini E, Hyttinen J, Paci M, Severi S. Simulation of the effects of extracellular calcium changes leads to a novel computational model of human ventricular action potential with a revised calcium handling. Front Physiol 2020;11:314

35. Plank G, Loewe A, Neic A, Augustin C, Huang Y-L, Gsell MA, Karablas E, Nothstein M, Prassl AJ, Sanchez J, Seemann G, Vigmond E. The openCARP simulation environment for cardiac electrophysiology. Comput Meth Prog Biomed 2021;208:106223

36. O’Hara T, Virág L, Varró A, Rudy Y. Simulation of the undiseased human cardiac ventricular action potential: model formulation and experimental validation. PLoS Comput Biol 2011;7:e1002061

37. Land S, Park-Holohan S-J, Smith NP, Dos Remedion CG, Kentish JC, Niederer SA. A model of cardiac contraction based on novel measurements of tension development in human cardiomyocytes. J Mol Cell Cardiol 2017;106:68–83

